# Spatial Suitability Modeling of Zoonosis: Implicated Risk Areas of *B. anthracis* and Trends under climate change Scenarios in Ethiopia

**DOI:** 10.1101/2020.11.27.400879

**Authors:** Michael A Yousuf, Solomon Asfaw, Shimelis Mengistu, Mohammedsham Husen

## Abstract

The causative agent of Anthrax *B. anthracis* has long been known to cause disease in animals and humans. Its worldwide distribution includes Ethiopia as an endemic country to the disease. The current study was aimed at identifying and developing risk maps, in areas that are suitable for the persistence of anthrax spores under climate change scenarios by using anthrax occurrence data and other predictor variables in MaxEnt model. A total of 158 occurrence locations were used as inputs along 10 current bioclimatic, future climatic grids and topographic covariates to develop a model and evaluate the individual contribution of each variable to the presence of *B. anthracis* in Ethiopia. It’s concluded that the most important variables limiting the distribution of *B. anthracis* in Ethiopia were Temperature, Precipitation, and Elevation. Under HADGEM2-ES future modeling scenarios except for RCP 8.5/2050 there is a decrease in areas of suitability from Current scenario under RCP 2.6/2050, RCP 2.6/2070, and RCP 8.5/2070. Subtle expansions of suitable areas are identified under RCP 2.6/2050 and RCP 2.6/2070 in the eastern parts of Ethiopia. However, there are small portions of southern areas that are expected to lose suitable habitats under all future scenarios. These findings could help health management authorities to formulate prevention and control strategies of anthrax in suitable areas under *B. anthracis.*

## Introduction

Modeling of suitable habitats for species distribution under climate change scenarios has received much attention in recent years due to its application in several fields of studies, which offer significant importance in associating the presence of the required species to its ecology. Recently, the application of these modeling techniques has gained more recognition in areas of Epidemiology, Conservation, Ecology, Evolution, Invasive species management, and other fields[1–2]. In epidemiology predictive modeling of disease-causing agents based on the underlying environmental variables of known occurrence sites to determine the possible risk areas of species and prognosticating its future distribution under climate change scenarios constitute the basic and major benefits of modeling techniques. As a result of widely recognized techniques to identify the potential and geographic areas of disease distribution in epidemiology, the modeling techniques can be applied in studying zoonotic diseases such as anthrax to determine its causative agent habitat suitability with climate changes.

Anthrax is an important global disease of Animals that carries high animal morbidity and mortality and is also associated with spillover to humans [3]. The disease is caused by gram-positive rod-shaped bacilli which belong to the family of *Bacillaceae* and called *B. anthracis* [4]. Usually, anthrax transmission to the animal is through ingestion of whether the spore and/or vegetative form of its causative agent from contaminated grazing land and for humans the transmission is either by contact, air, or ingestion; it's usually an occupational disease.

Anthrax distribution has got concerned globally, which is reported from all continents that are populated heavily with animals and humans [5]. Worldwide, an estimated 20,000 to 100,000 cases of anthrax occur annually, mostly in poor rural areas [6–7]. Due to a lack of proper control and prevention strategies and weak public health services, several poor African countries have often hyper epidemic/endemic status to the disease. For instance, Zimbabwe was the country of the largest outbreak of anthrax, with about 10,000 human cases from 1979 - 1985, concerning civil war which caused the interruption of public health services in the country [8].

Accordingly, Ethiopia is one of those African countries which report anthrax as an endemic disease. The disease occurs in May and June every year (anthrax season) in several farming localities of the country, although suspected cases of livestock anthrax are reported from several districts, few of those are officially confirmed [9]. The previous studies indicate that the disease is well recognized by rural communities but little is known about its prevalence, epidemiology, and public health significance.

The disease cycle of anthrax involves host species, predators, scavengers, insects, water, soil, and various environmental factors. The overall of the above requirements must be met for an anthrax outbreak to occur. Furthermore, anthrax disease is capable of maintaining its infectious spore banks in the soil for decades or even potentially centuries [10]. These complex ecological, epidemiological and microbiological mystery of *B. anthracis* necessitated multidisciplinary approaches to study the distribution of anthrax in the environment and spatial modeling techniques have been used as one of those approaches to determine *B. anthracis* habitat suitability [11–14]. However, it has been found that there is a little experience to extend the modeling of anthrax habitat suitability to the future climate change.

The usual way to determine anthrax habitat suitability and forecast its future distribution under climate change scenarios by modeling techniques is to identify and incorporate several environmental factors in the modeling that are known to influence its outbreaks in the environment. There has been extensive research regarding the suitability modeling of *B. anthracis.* For instance, [11] demonstrated that precipitation, vegetation, topography, and soil pH could influence the suitable habitats of *B. anthracis* and [12] established the influence of *B. anthracis* habitat suitability by precipitation, temperatures, and vegetation. However, although the effect of those factors in spatial modeling of *B. anthracis* was demonstrated in the previous years, a little attention has been paid in the extrapolation of those influential factors to determine spatial suitability modeling of *B. anthracis* under the future climate change scenarios.

The present article presents a set of selected environmental factors for modeling of *B. anthracis* spatial suitability on the current condition and extending the modeling to the future climate change scenarios. Based on these, MaxEnt modeling technique was employed to determine suitable habitats under both current and future scenarios.

## Materials and methods

### Study area

The study was conducted in Ethiopia, a country situated in horn of Africa; a land locked country that have an area of 1.104 million km^2^. Ethiopia’s national livestock population approximately consists of 57 million cattle, 30 million sheep and 23 million goats, and 57 million chicken, as well as camels, and equines. The highland parts of the country comprises most of these animals, where also the area that most of the country’s population live [15]. Ethiopia experiences a hot dry season between October and January and two rainy seasons; the periods of high rainfall which elapses from June to September, and relatively small rainy season which is between March and May “Fig 1”.

**Figure 1:**
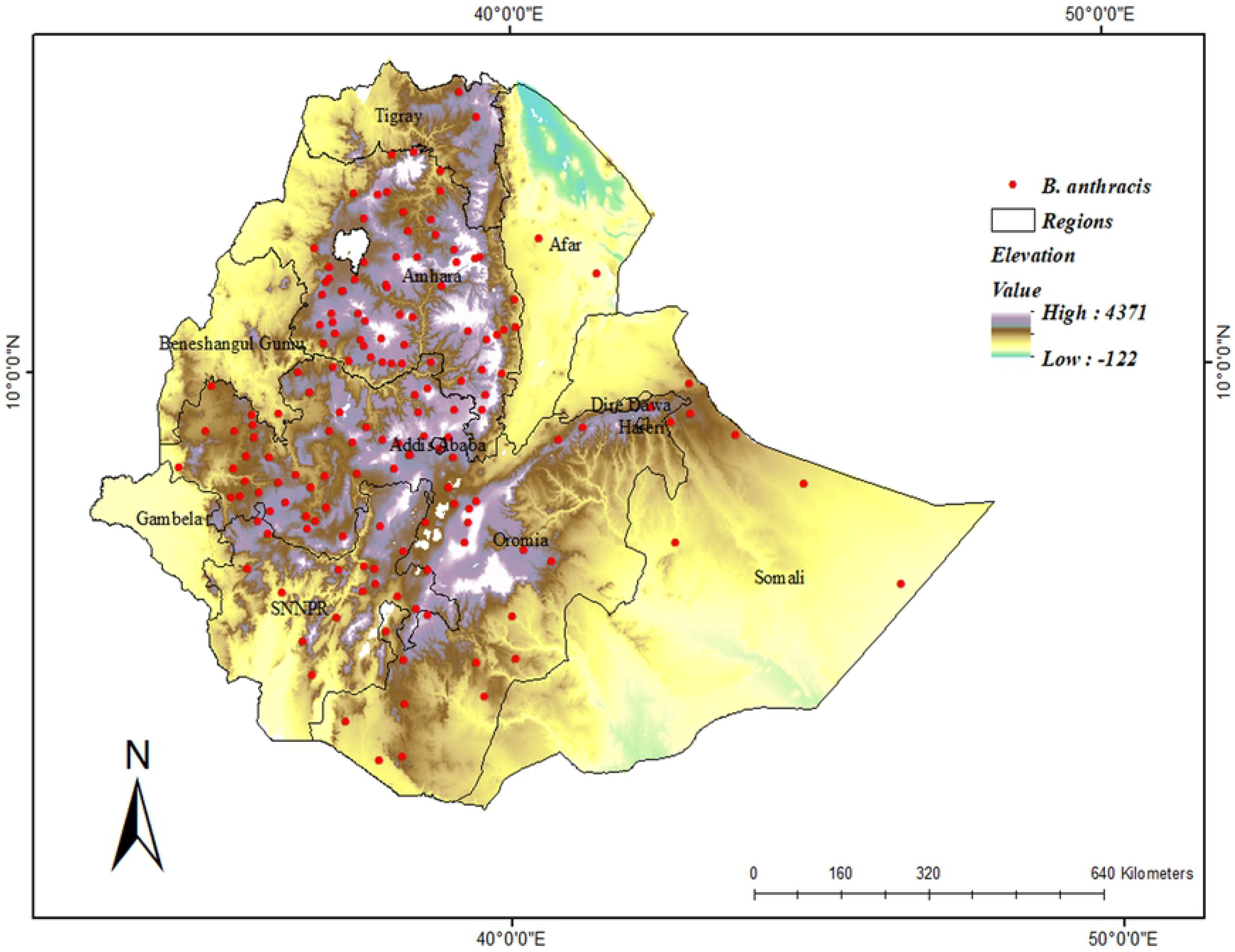
Map of Ethiopia depicting the country’s altitude, regions and *B. anthracis* occurrence points.

### *Bacillus anthracis* occurrence data

Anthrax cases for the years from 2014-2018 were the response variable of the model. The overall anthrax cases data in electronic format for the period of 2014-2018 were obtained from the Ethiopian Ministry Of Agriculture. Each anthrax case was indexed with their geographical location of the case, and characterized by region and Zones/woredas names in which the case presented. Those cases without coordinates, the reference points were given based on their areas of occurrence.

The dataset record includes a total of 7073 cases with a total death of 2286. Outbreak events in domesticated animals, large (cattle) and small (sheep and goats) ruminants, constituted the majority of the dataset. 158 total records were constructed from these case localities to develop a final model after the removal of duplicated cases and those without coordinate reference points from the dataset of 2014-2018.

Since the point data was in a spreadsheet format, it was converted to shapefile and used a spatial reference system of Universal Transverse Mercator (UTM), WGS84 Spheroid, and Datum with Zone 37N to use in ArcGIS.

The point data were standardized and converted to Universal Transverse Mercator (UTM) coordinates for use in ArcMap using Microsoft Excel. Only the species involved and UTM coordinates were used as input for the model. By using the Excel database a comma-separated values (.csv) file was created for use in MaxEnt.

### Predictor variables

The variables used as predictors were bioclimatic and topographic raster data. The Bioclimatic and Digital Elevation Model (DEM) data were downloaded from (www.worldclim.org) at 30 arc seconds resolution. To simulate the effect of climate change on the spatial distribution of *B. anthracis*, the study was used current bioclimatic and future scenarios climate grid data “Table 1”.

**Table 1:** Predictor variables used to model *B. anthracis* distribution in Ethiopia.

| Environmental variables | Codes | Source | Data type |
| --- | --- | --- | --- |
| Annual Mean Temperature | BIO1 | Worldclim | Climate |
| Temperature Annual Range | BIO7 | Worldclim | Climate |
| Mean Temperature of Wettest Quarter | BIO8 | Worldclim | Climate |
| Mean Temperature of Driest Quarter | BIO9 | Worldclim | Climate |
| Mean Temperature of Coldest Quarter | BIO11 | Worldclim | Climate |
| Annual Precipitation | BIO12 | Worldclim | Climate |
| Precipitation of wettest month | BIO13 | Worldclim | Climate |
| Precipitation of Driest Month | BIO14 | Worldclim | Climate |
| Precipitation of Wettest Quarter | BIO16 | Worldclim | Climate |
| Precipitation of Driest Quarter | BIO17 | Worldclim | Climate |
| Elevation | Alt | Worldclim | Elevation |

According to [16] a scenario is the consistent description of non-existed possible future events of the world. The Current climate events are represented by their averages for the period between 1950 and 2000 [17]. Future climate scenarios are represented by the projections for two time periods, 2050 (average climate conditions for 2041–2060), and 2070 (average climate conditions for 2061–2080) based on the IPCC fifth assessment report.

To model impact of climate change under low and high emissions scenarios on *B. anthracis* distribution in Ethiopia, the current study used the same 10 future climatic grids that were initially used for modeling of the current potential distribution of *B. anthracis* with about 30 arc seconds spatial resolution. The study was selected two future GHG (greenhouse gas) concentration trajectories, also known as Representative Carbon Pathways (RCP 2.6 and RCP 8.5), for two different periods (2050 and 2070) as adopted by the IPCC in its fifth Assessment Report (AR5) [16].

The first, RCP 2.6, is based on a reduction in Greenhouse Gas Concentration. According to this lowest emissions scenario, global annual Greenhouse Gas emissions peak between 2010 and 2020, and temperatures are projected to increase in range from 0.3 to 1.78°C. RCP 8.5 proposes an increase in Greenhouse Gas Concentration. Highest emissions assume Greenhouse Gas emissions will continue to increase throughout the 21st century, with temperature increasing from 1.4 to 4.88°C by 2100.

This study selected a global circulation model (GCM), HadGEM2-ES (Hadley Global Environment Model 2 Earth-System) developed by the Hadley Center, United Kingdom. HadGEM2-ES models have been used to perform ` all the CMIP5 (Coupled Model Intercomparison Project Phase 5) centennial experiments including ensembles of simulations of the RCPs. HadGEM2-ES is one of the models used by the international governmental panel on climate change (IPCC) in its fifth Assessment Report (AR5).

To identify the effect of climate change on *B. anthracis* potential distribution, this study developed an index reflecting the “suitability status change” by subtracting the climatic niche suitability values of the current events from the future conditions as suggested by [18]. As the values in each condition (current and future) are binary (presence: 1, and absence: 0), when these two values are added or subtracted, the only three possible results for each coordinate are a negative one (−1), a zero (0), and a one (1). These results were used as (−1) represents the no longer suitable area (currently suitable areas but that will not remain so in the future), (0) represents remaining an unsuitable area (unsuitable currently and under future climatic conditions), and a (1) represents remaining a suitable area (suitable currently and under future conditions).

### *B. anthracis* Distribution Modeling

Maximum Entropy (MaxEnt) modeling technique has long been used to model species distribution across the area of interest including disease causing agents and vectors [12–14] [19–20]. MaxEnt is the preferred method to model species distribution because of its simplicity to employ and can easily extrapolate the current distribution of species in charge into the future projectile climatic scenarios.

MaxEnt uses the presence only data and a number of environmentally gridded layers to produce a binary map of which its 0 value represent an area of low suitability to 1; high suitability area for predicted species. However, the use of presence only data in MaxEnt cannot purify the study from the sampling bias. Hence, mitigation of sampling bias was done by creating of the background layer in arcMap 10.3 to enable the selection of random sample from sampling distribution.

In addition to overcoming the problem of sampling bias from the occurrence points one modeler must beat the difficulties of model complexity that result in overly optimistic model outputs. MaxEnt have two built in parameters that control model complexity and fits (i.e. feature class and regularization multiplier), but their absolute performance is not in a default settings of the MaxEnt technique [21]. Therefore, it’s imperative to determine the best possible combination of features and regularization multiplier value before running MaxEnt model to generate the final representative model. In doing these, first the highly related collinear environmental variables are tested and removed from the predictor covariates.

Accordingly, variables with a VIF < 10 were retained. After the removal of collinear predictors, the remained 7 variables were used to simulate the process of feature selection and to determine the value of regularization multiplier. The ***usdm*** package of R employ VIF to detect the collinearity among the predictor variables in which the VIF threshold greater than 10 indicates the collinearity of that variable to others, this means there is a correlation coefficient close to 1[22].

Premodelling MaxEnt model setting selection preference was carried out by using ENMeval package of R Statistical Software by using variables that have no collinear problems to limit the model Complexity from the modelling procedure [23]. Thus, the ENMevaluate result of ΔAICc (delta akaikae information criterion adjusted for small sample size) which is equal to 0 was chosen. ΔAICc has been used by several authors to select the best possible features of environmental layers and to choose the corresponding Regularization multiplier prior to running MaxEnt modeling [19–20] [24].

The number of replicate runs was set to 10, and replicate run type to Cross-validate. Cross-validation has one big advantage over using a single training/test split: it uses all of the data for validation, thus making better use of small data sets [25]. The advanced options in MaxEnt that were selected included the maximum iteration set to 5000 to allow the models to have enough time to reach convergence at 0.00001[26]. 90% sensitivity was set within the MaxEnt model for determining suitability.

### Model evaluation

The area under the Receiver Operating Characteristics (ROC) was used to assess the accuracy of the model in cross-validation. In the MaxEnt model, the AUC of the receiver operating characteristic plot is used as a default evaluation criterion to assess the accuracy of the model [27]. To provide information about the degree of contribution of each variable, response curves and table of percentage contribution of each variable were used. The jackknife test of variable importance was also used to measure the model performance of our modeling results. The overall process of modeling summarized in “Fig 2”.

**Figure 2:**
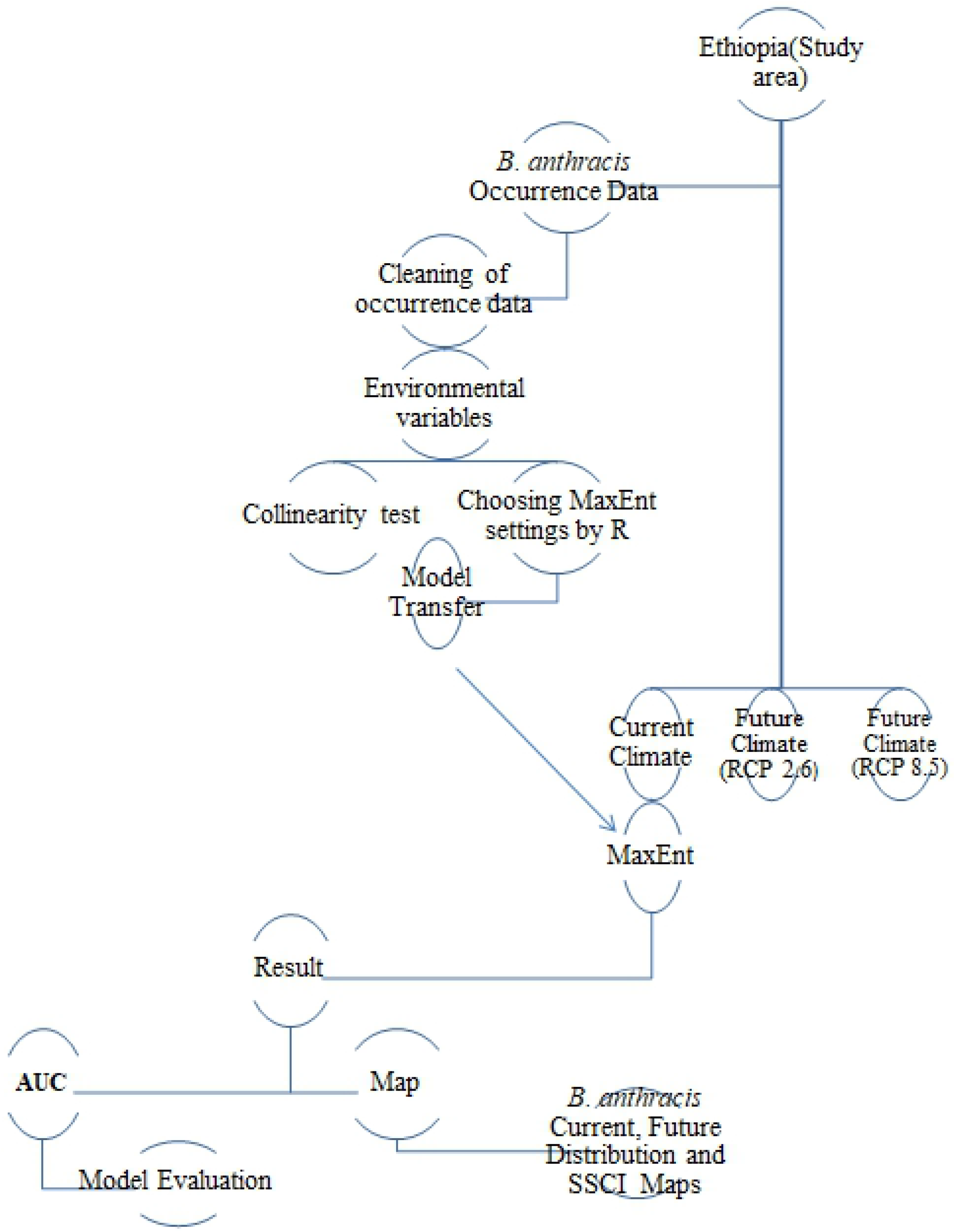
Flow-diagram summary of the study indicating the modeling process, evaluation and development of suitability maps.

## Results

### Accuracy metric

The Modeling processes for the current *B. anthracis* distribution and each of the four Future scenarios had reached the convergence of accuracy (0.0001) before the maximum iteration setting of 5000. The MaxEnt model had a regularization multiplier of 4, allowed the Linear (L), Hinge (H), Product (P), and Quadratic (Q) features, The ROC curve had an average AUC of 0.75 and was significantly different from a line of no information (p < 0.01) “Fig 3”.

**Figure 3:**
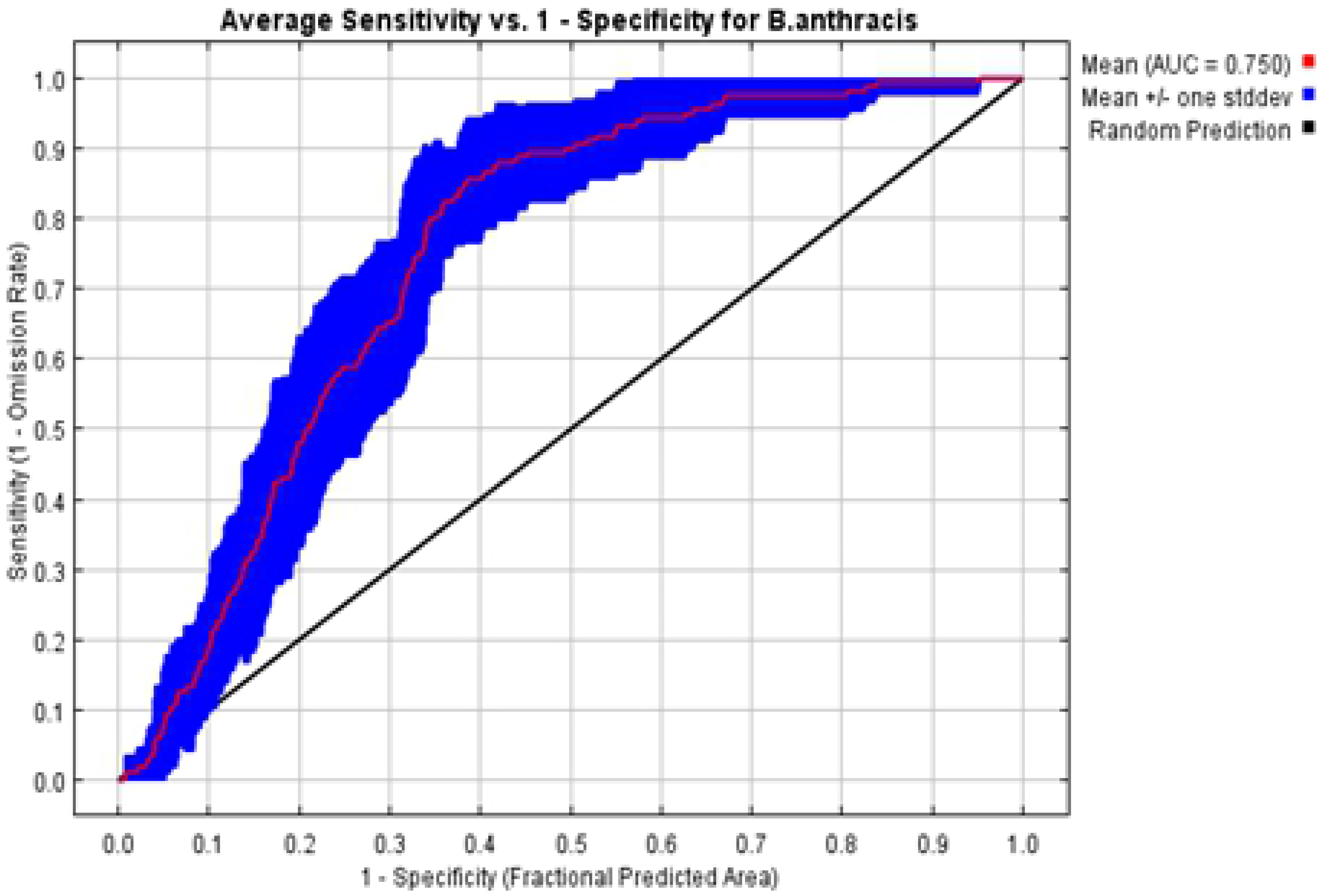
The average test of AUC for the 10 model replication runs for *B. anthracis* distribution.

Out of 11 selected predictor variables, 4 of them are excluded as they have a collinearity problem, i.e. Bio9 (Mean Temperature of Driest Quarter), Bio11 (Mean Temperature of Coldest Quarter), Bio14 (Precipitation of Driest Month), and Bio16 (Precipitation of Wettest Quarter). The remained environmental variables are contributed to the model significantly.

Looking at the estimation of the contribution of each predictor variable to the model, the table of percentage contribution and permutation importance showed that after Annual Mean Temperature, at 50.7%, elevation (23%), Mean Temperature of Wettest Quarter (9.7%), and Temperature Annual Range (9.4%) contributed most to the model “Table 2”. The other remained variables contributed relatively little.

**Table 2:**
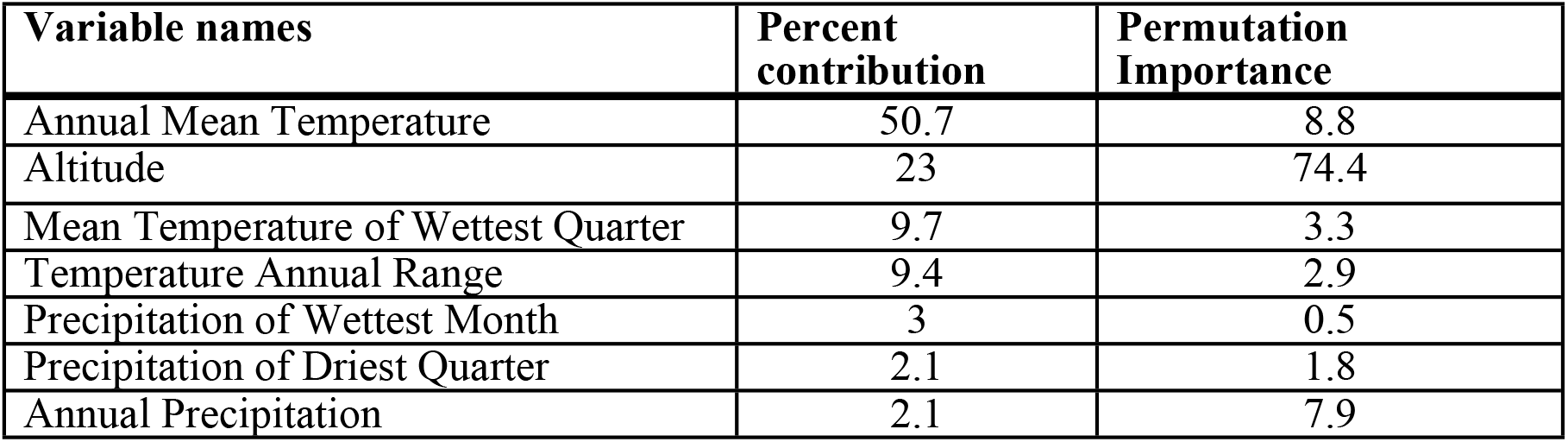
Table indicating Variable contribution includes percent contribution and permutation importance of each variable.

### Variable importance

The jackknife of AUC for *B. anthracis* distribution indicated that Mean Temperature of Wettest Quarter was the most important contributing variable to the model, followed by Elevation and Annual Mean Temperature. However, the variable that decreased the gain the most when omitted was the Annual Mean Temperature, which means that it had the most information that was not present in other variables “Fig 4”.

**Figure 4:**
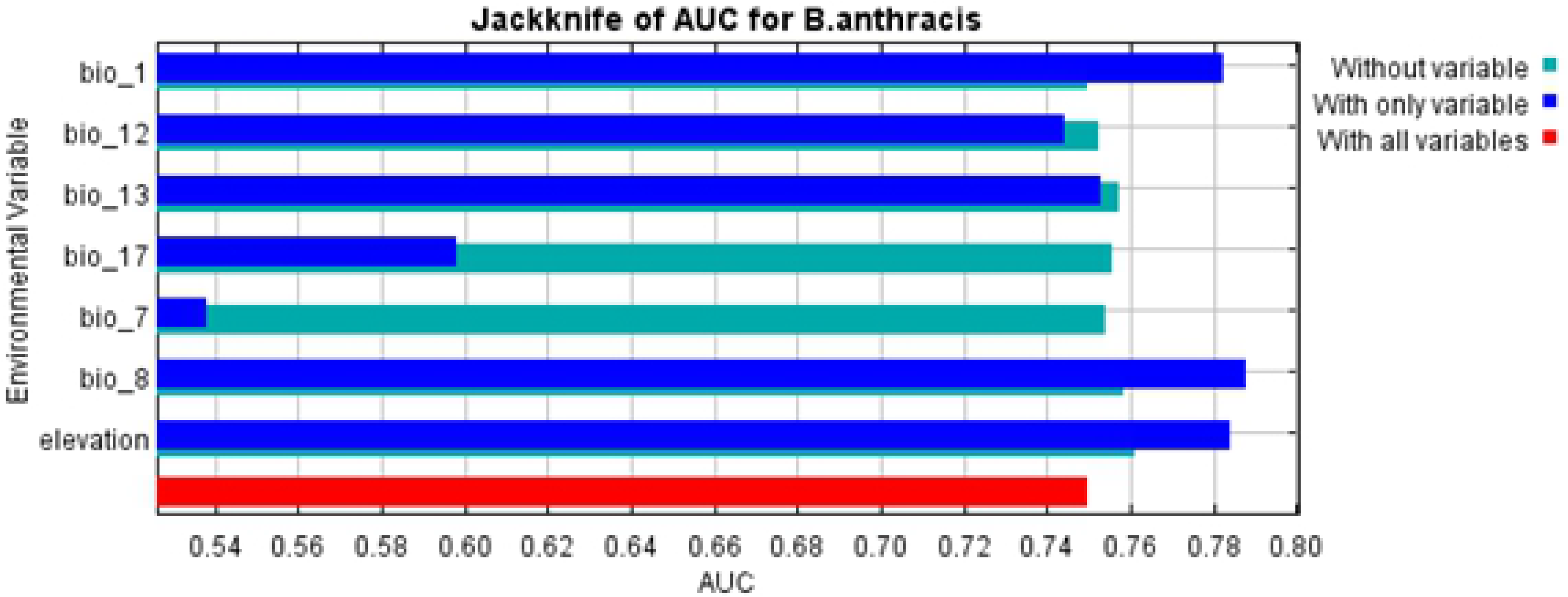
Jackknife of AUC result for Variables used to construct the MaxEnt Model.

Looking at the AUC of the Jackknife AUC, the most significant variables with scores of > 0.75 were Elevation, Annual Mean Temperature, Mean Temperature of Wettest Quarter, and Precipitation of Wettest Month. See Response curves for these variables about their suitability for the prediction of *B. anthracis* spore potential distribution in Ethiopia.

MaxEnt model response curves showed how the model’s predictions changed as the environmental variables varied. The probability of *B. anthracis* presence increased as the values of elevation increased up to 3000m then declined slightly as elevation values increased above“S1 Fig”.

The Response curve of *B. anthracis* in relation to the Precipitation of the Wettest Month represents how Habitat suitability changes with changes in the Precipitation of the Wettest Month. The x-axis shows the Precipitation of the Wettest Month while the y-axis is the Probability that the habitat is suitable for *B.anthracis*. The figure shows that Habitat suitability of *B.anthracis* slowly increasing from 0 units of Precipitation of the Wettest Month, and become constant above the 400 units of Precipitation of Wettest Month.

### Current and future scenario of Suitability maps for *B. anthracis*

The Current scenario model identified the following regions as suitable for the Persistence and Distribution of *B. anthracis* in Ethiopia. These areas include Eastern, Southern and Western Oromia Region, Eastern Afar region bordering Eritrea, Northeastern quarter of Somali Region, Northern and Central SNNPR and the Northern Tier of the Country in which the Suitable area stretches from the Topmost Northern parts of Tigray to the Central area of Amhara Region surrounding Lake Tana where the predictions then extend to the Southward of Northern Shoa Zone “Fig 5”.

**Figure 5:**
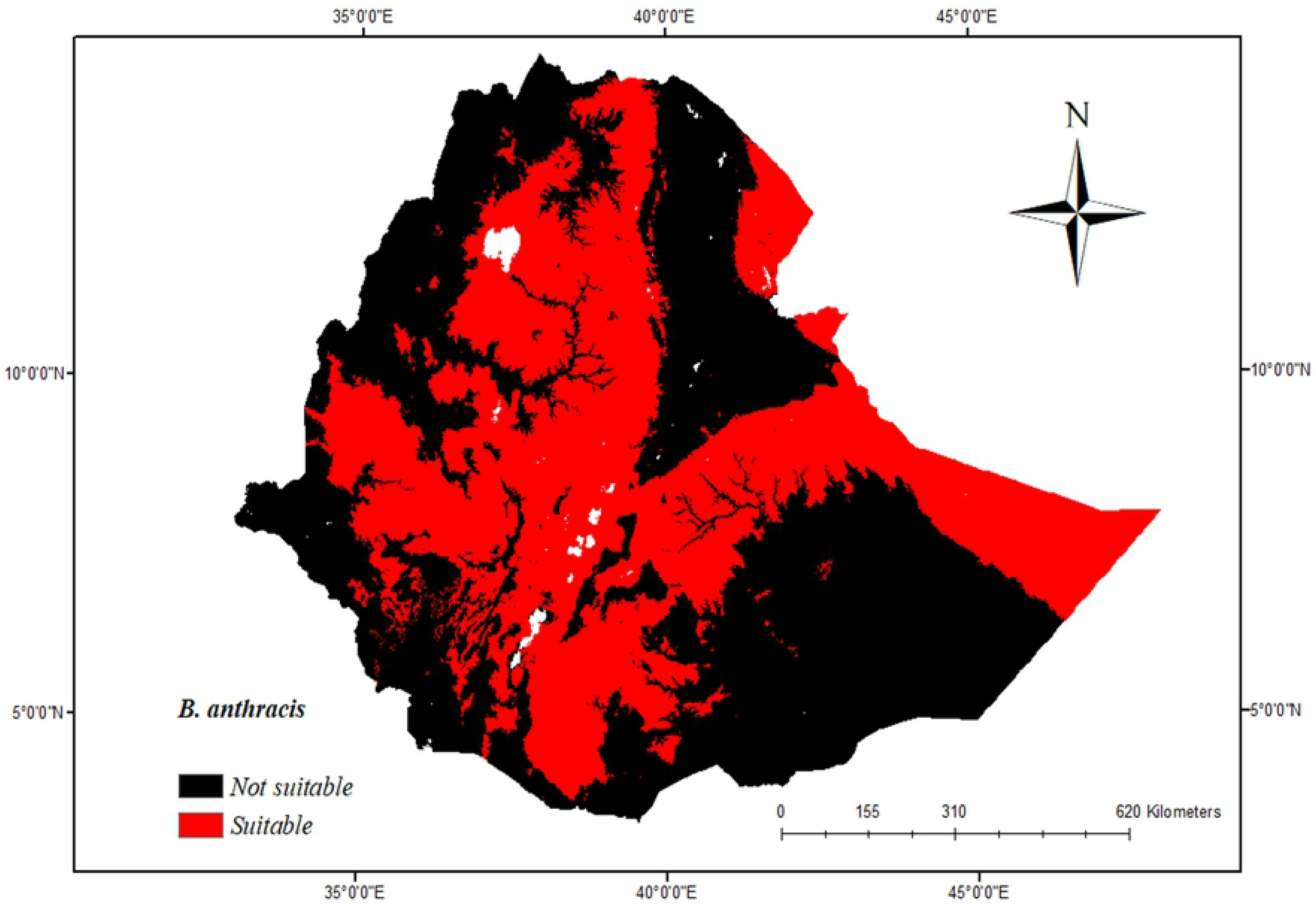
Binary Map of *B. anthracis* Suitable areas in Ethiopia under the current bioclimatic scenario.

For the four future models, the AUC results were very similar. This similarity is to be expected with similar data. The results of four future models are graphically similar in appearance to Figure 4. Results for all four future models closely resembled the variable importance for the Current scenario model. Mean Temperature of Wettest Quarter is always the highest contribution, followed by Elevation.

Under the optimistic scenario of RCP 2.6/2050, there is a decrease in suitable areas of *B. anthracis* even if the occupations are clearly observed in the eastern parts of Ethiopia. A substantial habitat loss was observed in the eastern Somali areas of Dolo and Jarar bordering Somalia and the some eastern parts of Afar region bordering Eritrea. Contraction of suitable areas also occurred in some western parts of Oromia region bordering Sudan and Southern Omo parts of SNNPR bordering Kenya. The recession of habitat suitability also occurred in Northern Waghimra and Gondar areas of Amhara, the Northernmost, and some central parts of SNNPR and some Eastern parts of Tigray. Complete loss of Habitat for *B. anthracis* was observed in the Bench-Maji and Southern Omo Zone of SNNPR while the remaining parts of the Country retain a suitable Environment for *B. anthracis*.

The increased concentration of Greenhouse Gases under RCP 8.5/2050 will increase the *B. anthracis* habitat suitability area in the year of 2050, in the southern Bale which stretches toward the Borena parts of the Oromia region and center of the study area including the most highland parts of the Country. There are also several very small areas of expanded habitat scattered across the Eastern Ethiopia in Afar region.

Considering the RCP 2.6/ 2070, No geographic shifts and habitat loss were evident compared to RCP 2.6/ 2050 except a subtle decrease in habitat expansion in the most eastern parts of Somali region around the southern areas of Jarar and Dolo, while most parts of the remaining region will remain suitable for Persistence of *B. anthracis* spores across the country.

Under RCP 8.5/2070, the model suggests a huge loss of habitat suitability for anthrax spores in 2070 in which the geographical distribution is limited to the central and northern highland parts of the Amhara region. A noticeable change will occur in eastern and southern parts of the Oromia region including western areas where a suitable environment will become narrowly connected exhibits a little contracting suitable climate for *B. anthracis.* The overall anthrax predicted binary maps of RCP scenarios are shown in “Fig 6”.

**Figure 6:**
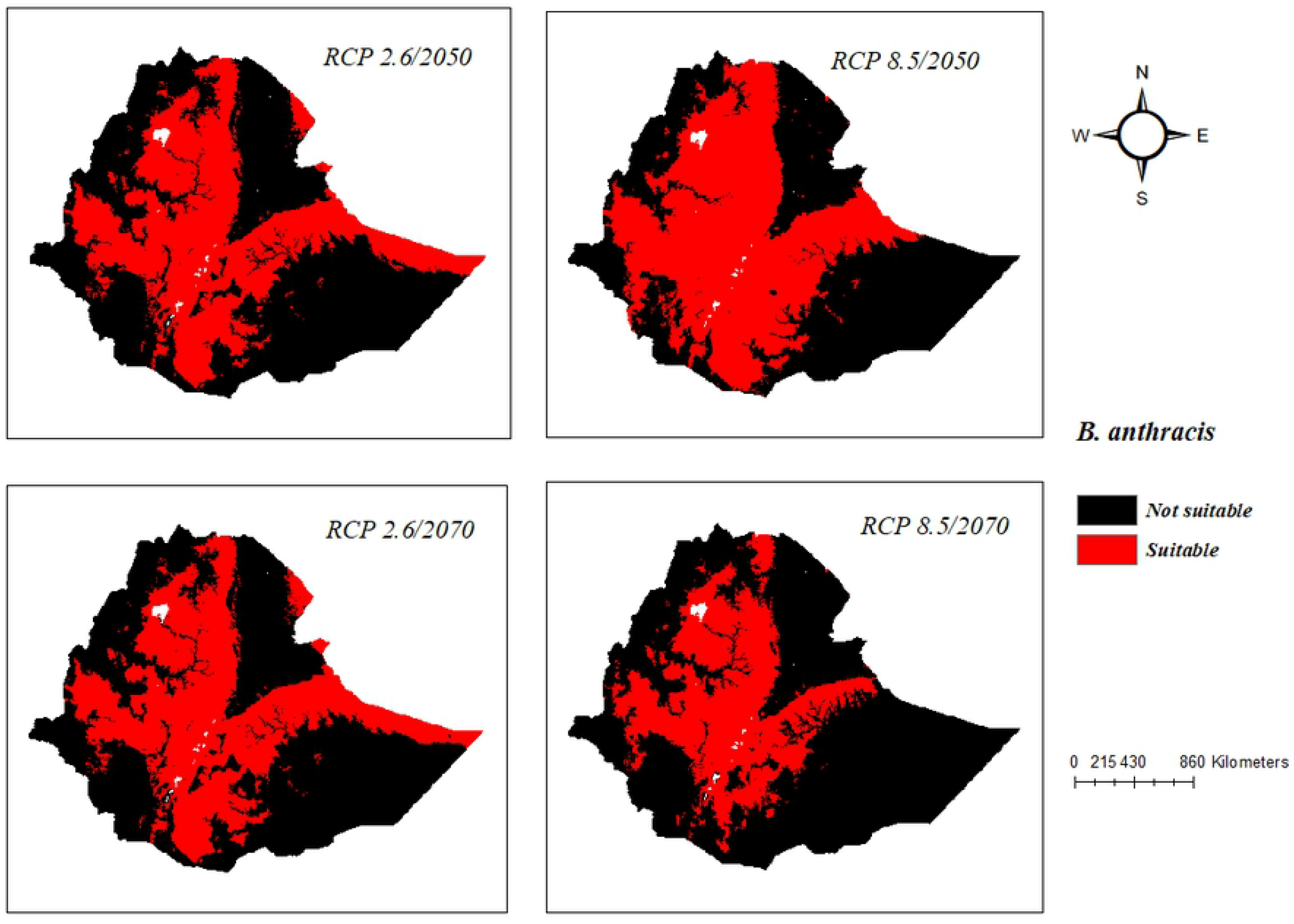
Future potential distribution of *B. anthracis* predicted by the HADGEM2SE model under RCP 2.6 and RCP 8.5 scenarios.

### Suitability Status Change Index

A comparison of suitability differences between the current and future scenarios was carried out by using the “Suitability status change index” as suggested by [18]. When the areas between Current and Future conditions are compared, differences between the models are evident (S2 Fig). These differences in area are summarized in “Table 3”.

**Table 3:** Suitability status change index area for future climate change scenarios.

| Suitability area* | 2050 |  | 2070 |  |
| --- | --- | --- | --- | --- |
|  | RCP 2.6 | RCP 8.5 | RCP 2.6 | RCP 8.5 |
| Remain suitable area | 3902 | 104555 | 4970 | 10712 |
| No longer suitable | 76620 | 95049 | 76071 | 213546 |
| Remain unsuitable | 104493 | 925855 | 104441 | 901201 |

\* Area measured in km<sup>2</sup>

## Discussion

In this study, Bioclimatic and DEM variables were used to predict the current and future spatial suitability of *B. anthracis* in Ethiopia. The models proved to be statistically significant with an AUC of 0.75 for both Current and Future Suitability scenarios.

This study identified that a narrow band of Temperatures, Precipitations, and Elevation can limit the distribution and persistence of anthrax spores in Ethiopia. Ethiopia is a country which has diverse climatic condition due to the situation of its topography. Altitude is the second most important to determine the suitability of the environment for *B. anthracis* spores. This finding is in agreement with the result of [12], who defines the anthrax belt in the contiguous USA by specific bands of NDVI, precipitation, and elevation.

The study found that temperature abnormalities can describe the distribution of *B. anthracis* across the country. Mean Temperature is the most important variable to define *B. anthracis* distribution in Ethiopia. The recent climate condition of the country indicates that the Mean annual temperature increased by 1.3°C between 1960 and 2006, an average rate of 0.28°C per decade [28]. The increase in temperature can result in the variability of rainfall throughout the country which causes heavy flooding due to global warming [29]. This condition could create favorable environments in which the transmission and persistence of anthrax spores may maintain itself due to an increase in Mean Temperature along with the combination of Precipitation of the Driest Month in Ethiopia [9].

Although there were differences in results among the four climate models for both 2050 and 2070, all predictions entailed a significant reduction of the present-day distribution of suitable climatic conditions for *B. anthracis*, except under the projection of RCP 8.5/ 2050 in which a subtle increase of suitable area from current distribution is identified. Under the RCP 8.5/ 2050 model scenario the overall high potential distribution of *B. anthracis* habitat increased by 84497 km^2^ across the study area that may be due to changes in climate that mostly hit the western and central northern parts of Ethiopia.

Suitability is projected to decline in the southern parts of the SNNPR and Somali regions under the RCP 2.6 of both 2050 and 2070. The decrease in projection in the southern parts of Ethiopia is maybe from one of the following factors which might be due to other non-bioclimatic variables that are not used in this study under climate change scenario to limit the *B. anthracis* from further expansions across the large areas of southern latitudes or the response of decreasing suitable habitats which consist of grassland and pasture areas used for grazing activities of livestock and simultaneously rangeland expansion to the central and northern parts of the country which might lead to the migration of livestock upward to the northern latitudes because of changes in temperatures and precipitation.

These results are partially in geographical agreement with that of [30] who described the similar results of southern parts contraction in a suitable environment for *B. anthracis* in Kazakhstan under multiple climate change scenarios. However, the predicted trends of these changes are not as violent as that Kazakhstan. [31], demonstrated the effect of climate changes on the geographical distribution of *B. anthracis*, and found that the contraction of southern areas of suitable habitats than Northern latitudes of USA.

Other bacterial zoonotic study is also indicating contraction of the southern latitudes due to climate changes in North America. [32] Predicted similar contraction of southern latitudes in the study carried out to determine the effects of climate change on the distribution of plague and tularemia in USA. However, the propositions of contraction of the southern area distributions of these findings are not well understood. The conclusion based on the continental comparison might be warranted as the future climatic conditions of Northern America, Asia’s and Africa may differ based on several multi environmental factors.

In contrast to the above three scenarios, under scenario RCP 8.5/2070, the highly suitable areas decreased from the present until 2100. This indicated that different emission scenarios have different and opposite effects on the potential distribution of *B. anthracis* in Ethiopia. Relative to habitat gain and loss of *B. anthracis* under different climate change scenarios in Ethiopia, this study found that unchanged or little decreasing suitability of the environment to the northern latitudes of the country.

It has been described that the influence of climate change will decrease the livestock population in arid and semi-arid regions of Ethiopia. The trends of 2050 high emission scenario identified that the lowering of animal production in lowland parts of the country [29]. These finding is in line of our predictions in which the lowland areas of Somali and Afar regions loss the suitable habitats of *B. anthracis* under RCP 8.5 of both 2050 and 2070. This prediction might be indicative for anthrax distribution based on the availability of vegetation and rainfall that could determine the presence and migration of livestock from low suitable area to high suitable area of *B. anthracis* and in opposite direction.

## Conclusions

The current study is the first to apply HADGEM-2SE future climate models to map the distribution of *B. anthracis* in Ethiopia. Similar future predictions of anthrax spores build upon the process of this study should be carried out repeatedly to ensure the accurate predictions of forecasted changes in habitat suitability of *B. anthracis* and it’s also strongly recommended. Hence, this paper may provide a strong foundation for future research in modeling of suitable habitats for *B. anthracis* by using several other modeling techniques including MaxEnt which may subsequently provide more information to both animal and public health officials about the disease distribution. Depending upon the result of this study, Animal health protection officials and public health authorities could be benefited from it in formulating the protection, prevention, and control policies and strategies for suitable areas of *B. anthracis*.

## Supporting information

**S1 Fig: Response curves**

**S2 Fig: Maps of suitability status change for future distribution of *B. anthracis***

